## Supplementary information for "Molecular mechanism of proton-coupled ligand translocation by the bacterial efflux pump EmrE"

Movie S1. The dominant ligand transport pathway, as postulated in Fig. 2. After entering the binding site, the ligand accelerates the conformational change that subsequently allows for preferential protonation of E14$^B$. A second protonation event allows for rapid escape of the ligand, leading to the apo state.

Movie S2. Water wires and desolvation in the apo state. The inset shows Gaussian-smoothed probability of Grotthus wire formation with the top (orange) or bottom (blue) compartment as a function of time. After ca. 1 microsecond, the rearrangement of M21 and W63 creates a watertight seal that disrupts wires extending to the binding site, thereby preventing futile proton transfer.

Movie S3. Spontaneous ligand entry (left) and escape (right) trajectories obtained from Westpa, along with independently calculated free energy profiles for the fully deprotonated (orange) and doubly protonated (blue) systems. Residues that come into contact with the P4P ligand are shown in yellow.

| pK$_a$ values | | |
|---|---|---|
| | 1A Alchemical | 1B Alchemical |
| Ligand form | 6.57 | 8.9 |
| Apo form | 7.15 | 7.6 |

Table S1: The pK$_a$s obtained from the alchemical approach (described in Methods).

ΔpK$_a$s were calculated from the Henderson-Hasselbach equation using ΔG values obtained with Bennett Acceptance Ratio (BAR), and added to the reference value for free glutamic acid in solution (4.25).

$$\Delta G = -RTlnK_a$$
$$\Delta pKa = \frac{\Delta G}{2.303RT}$$
$$pKa_1 = pKa_2 + \frac{\Delta G}{2.303RT}$$
$$pKa_1 = pKa_2 + 0.174\Delta\Delta G$$

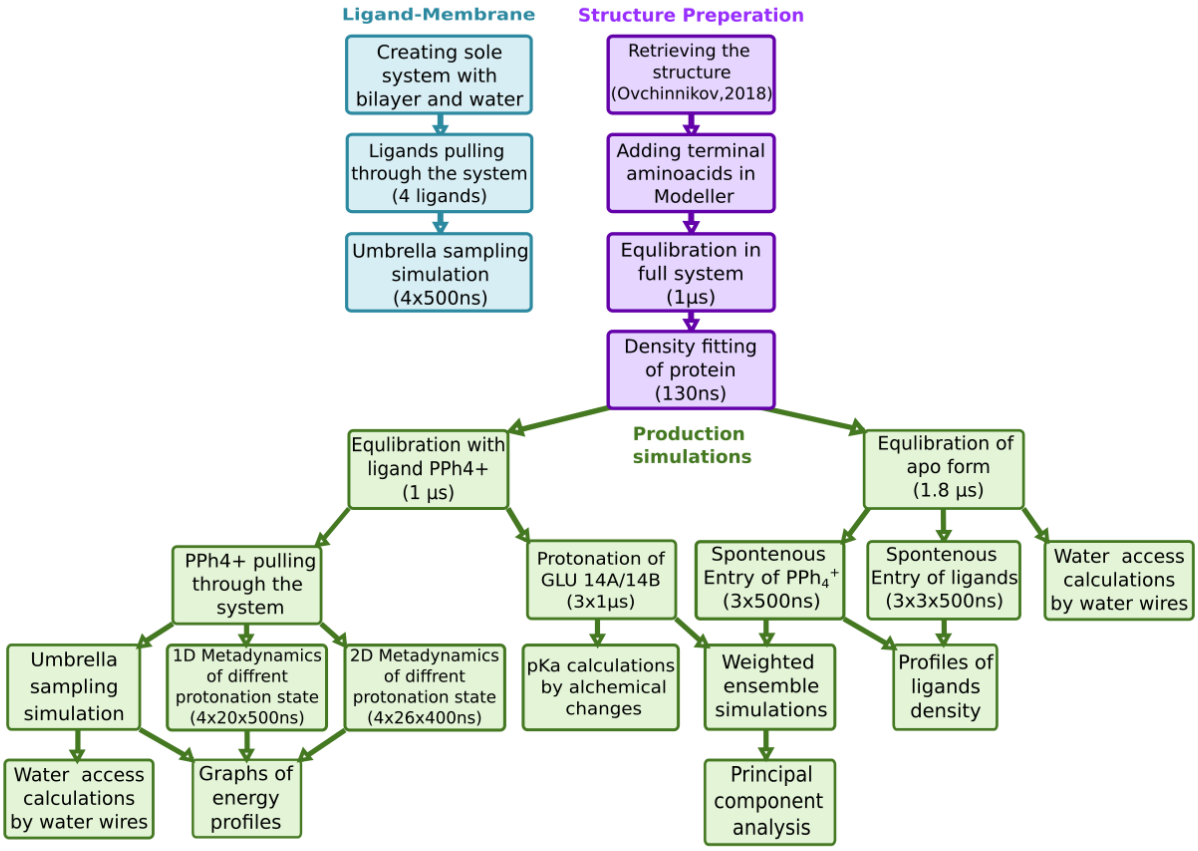


Figure S1: Workflow of the refinements and simulations performed in this study. See Methods for the details of all approaches used.

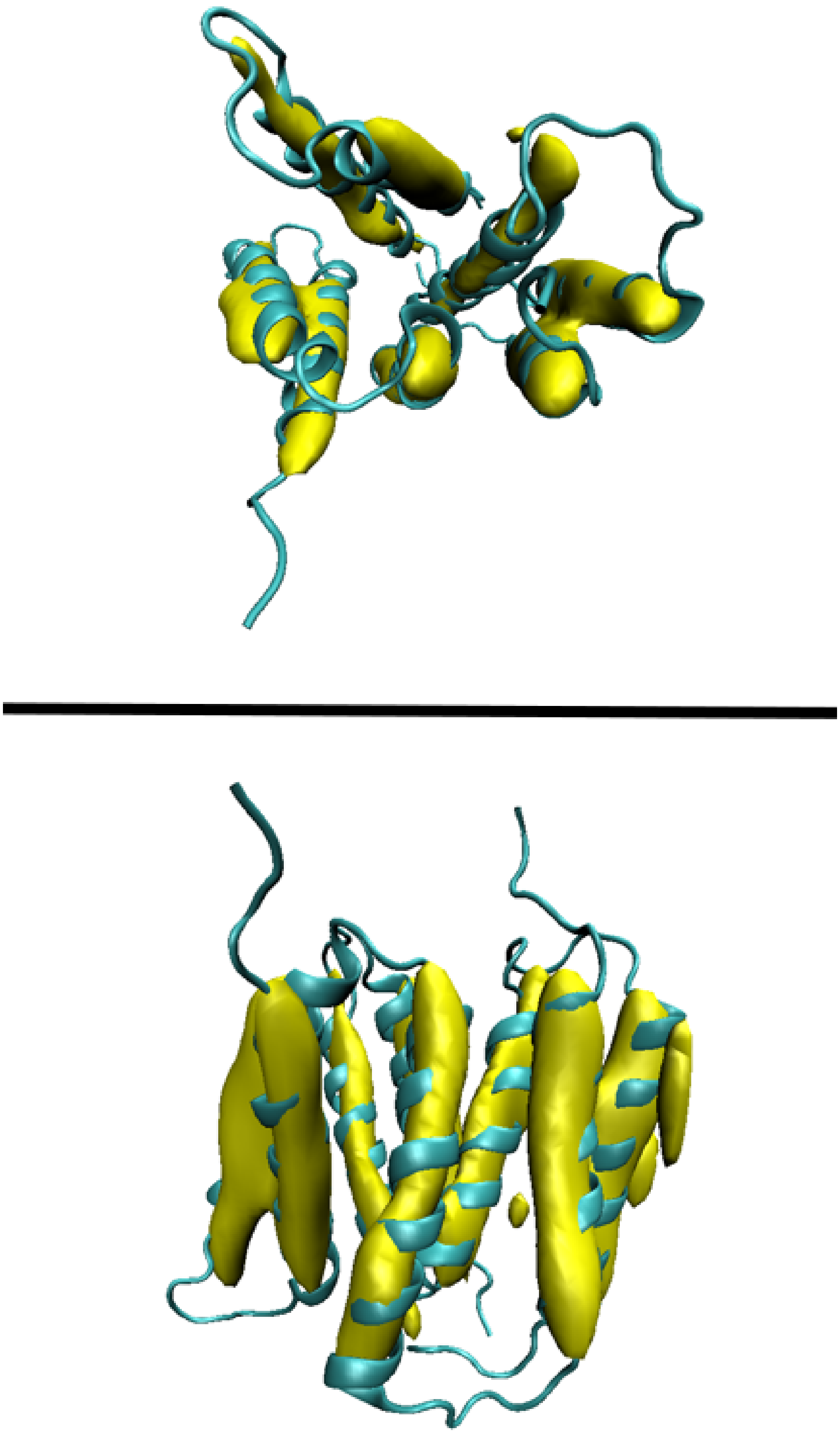

Figure S2: Alignment of the refined structure with the cryo-EM densities. Using an early implementation of the cryo-EM module in the Gromacs package, we set the densfit-sigma parameter to 0.45 nm and densfit-k changed linearly from 0 to 10000 over 5 ns and remained at 10000 for the subsequent 95 ns. In the implementation, the "forward" model of the electron density is created by Gaussian-smoothing the atomic structure, and the forces are then applied to minimize the cross correlation between the simulated and actual density. This method allowed to obtain a structure with a proper spacing and orientation of helices, corresponding to the PDB entry 3B5D.

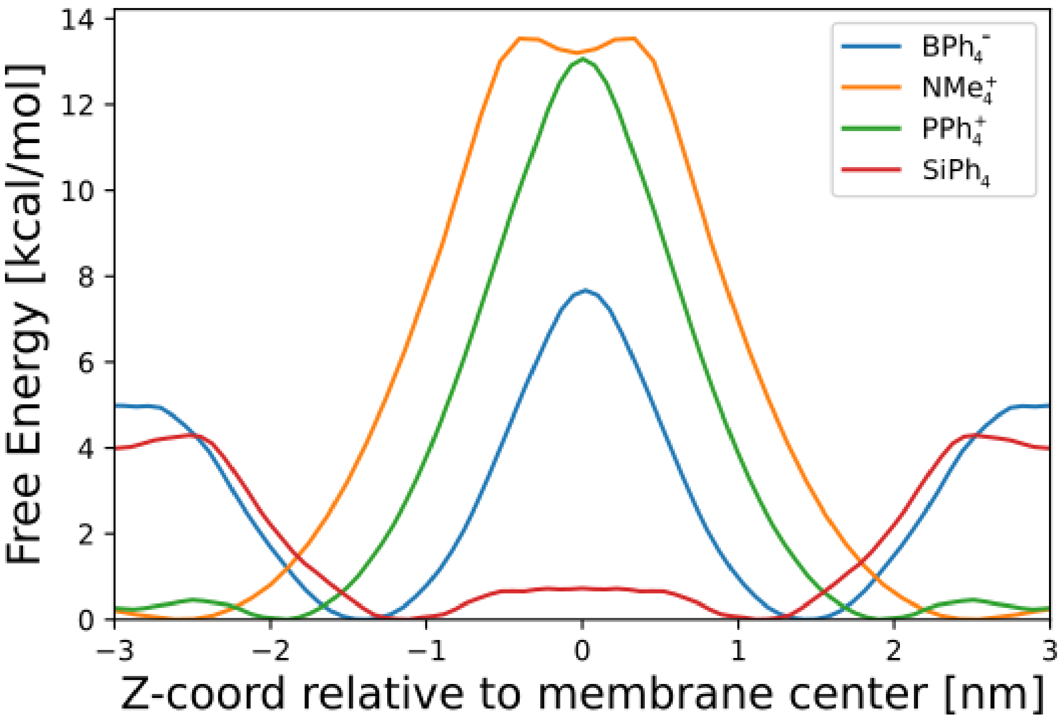


Figure S3: Free energy profiles for a single ligand passing through a pure POPE/POPG membrane. The membrane was generated using the CHARMM-GUI webserver. The umbrella sampling protocol was applied to four different ligands (aromatic: cationic $PPh_4^+$, neutral $SiPh_4$, anionic $BPh_4^-$, and non-aromatic hydrophobic cation $NMe_4^+$) to create free energy profiles as a function of the z coordinate. The profiles indicate that charged ligands preferentially reside in the aqueous phase near the bilayer surface, the anionic ligand resides in the headgroup region and can likely pass through the membrane in a spontaneous manner, while the neutral tetraphenylsilane shows sizeable affinity for the bilayer interior.

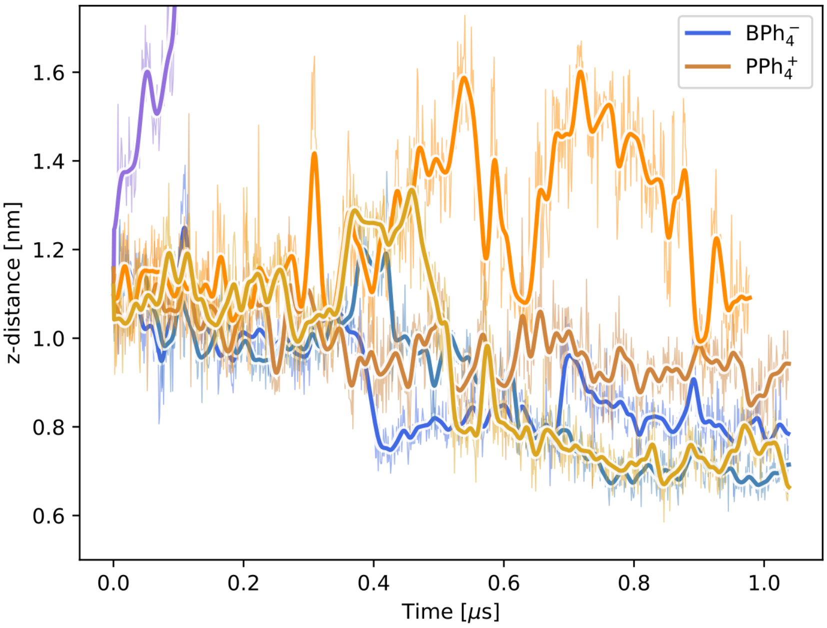


Figure S4: The relative z-coordinates of ligands during the 1-s charge-swap simulations. 3-channel ligand structures (distance from to the binding site between 1.0 and 1.2 nm) were taken from the equilibrium simulation and six simulations of 1  s each were performed for 3 systems with $PPh_4^+$, and 3 for $BPh_4^-$. The performed simulations were run as "as it is" or by changing the charges while maintaining the geometry. Of the 3 systems containing $PPh_4^+$ (yellow / orange / brown lines), all remained stably related to the channel entry, mainly going deeper (up to 0.8   nm) into the channel. At the same time, the least bound $BPh_4^-$ dissociated in one of the systems (purple line), while the other two progressed deeper into the binding channel, as did their cationic counterparts (blue / turquoise lines)

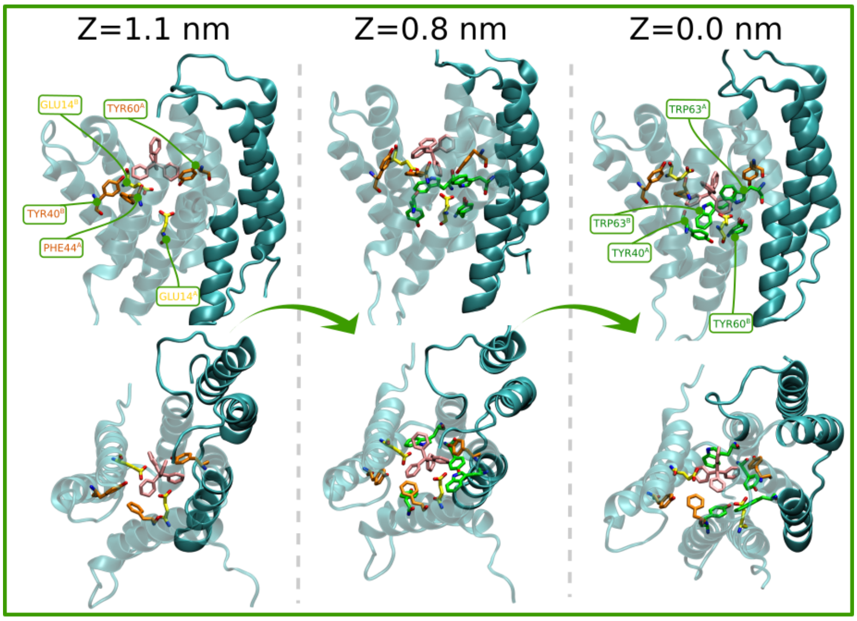


Figure S5: Visualization of the spontaneous ligand entry into the active site. Simulations obtained from Westpa provide an insight into the key amino acids involved in the ligand entrance into the binding site, here visualized at three crucial stages (Z = 1.1 / 0.8 / 0.0) where the ligand dwells due to the presence of free energy barriers. The amino acids actively involved in the transport mechanism are shown: E14, Y60, Y40, F44 and W63.

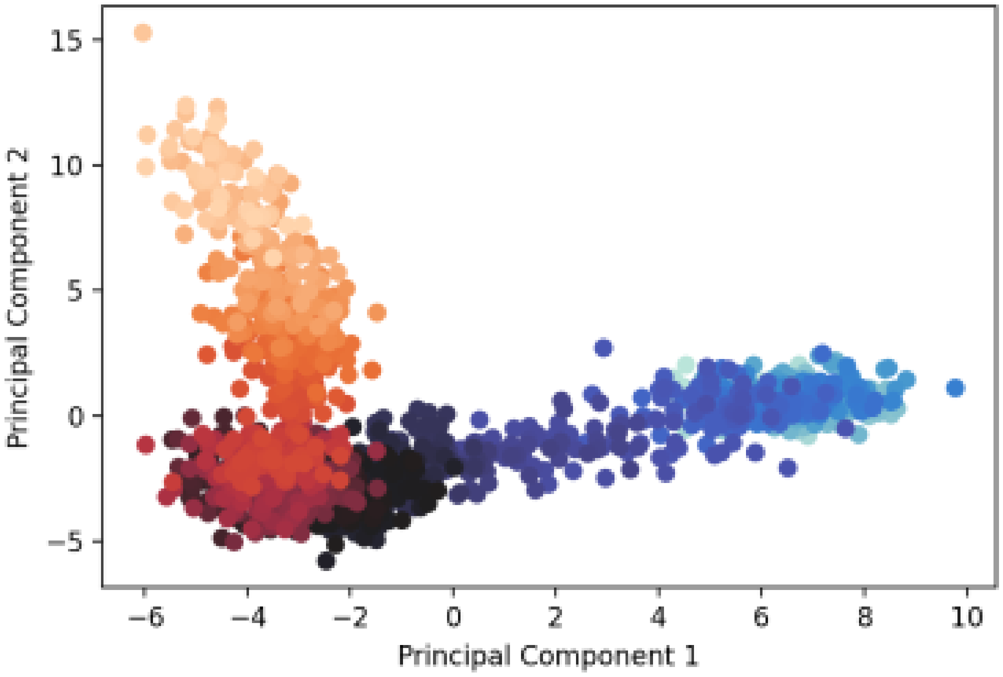


Figure S6: Projection of the spontaneous ligand exit trajectory on the two main principal components describing correlated changes in H-bonding patterns. The short trajectory from Westpa was merged with a follow-up 100 ns equilibration.
